# Cell Cycle Dependent Orchestration of Surface Layer Biogenesis in *Caulobacter crescentus*

**DOI:** 10.1101/2023.06.14.544926

**Authors:** Matthew Herdman, Andriko von Kügelgen, Ulrike Schulze, Alan Wainman, Tanmay A.M. Bharat

## Abstract

Surface layers (S-layers) are proteinaceous, two-dimensional crystals that constitute the outermost components of many prokaryotic cell envelopes. In this study, we investigated principles of S-layer biogenesis on the outer membrane in the bacterial model organism *Caulobacter crescentus*. Fluorescent microscopy revealed localised incorporation of new S-layer at the poles and mid-cell, consistent with elongation and division phases of the cell cycle. Next, light microscopy and electron cryotomography investigations of drug-treated bacteria revealed that bacterial actin homologue MreB is crucial for localised S-layer insertion. We further uncovered that S-layer biogenesis follows new peptidoglycan synthesis and localises to regions of high cell wall turnover. Finally, correlated cryo-light microscopy and electron cryotomographic analysis of regions of S-layer insertion showed the presence of gaps in the hexagonal S-layer lattice, contrasting with other S-layers completed by defined symmetric defects. Our findings provide insight into how *C. crescentus* cells form an ordered S-layer on their surface, providing evidence for coordination between the biogenesis of the cell envelope at multiple levels.

## INTRODUCTION

Cell envelopes of prokaryotes are complex, multi-layered structures that fulfil a variety of roles, such as mediating interactions with the environment including neighbouring cells, regulating import and export of material, and protection from external attack^1, 2^. Many prokaryotes including archaea, Gram-negative and Gram-positive bacteria express a macromolecular, proteinaceous sheath known as the surface layer (S-layer) as the most exterior part of their cell envelope^3–6^. There is increasing evidence suggesting that S-layers are abundant in prokaryotes, with a majority of bacteria and most archaea expressing an S-layer on their envelopes^4, 7^. S-layers are two-dimensional lattices made up of repeating copies of S-layer proteins (SLPs). Since SLPs are the highest copy number proteins in many prokaryotic cells, by many estimations they are one of the most abundant protein family found in nature^3, 8, 9^. Given their position as the outermost component of the cell envelope, it is no surprise that S-layers are important for several aspects of cell biology and are suggested to be an ancient form of a cellular exoskeleton^10, 11^. S-layers have been implicated in maintenance of cell size and shape^11, 12^, evasion from predators^13^, attachment to substrates^14, 15^, and as a protection against a range of environmental pressures^16–18^.

The understanding of the evolution of S-layers is far from complete, since SLPs appear in even the most ancient lineages of prokaryotes and show a high level of sequence variability^4, 12, 19, 20^. Despite this variability, S-layers share several organisational features; for example SLPs are often bipartite in nature, encoding distinct lattice-forming and cell-anchoring domains, often within the same protein^4, 21–23^. Secondly, many SLPs utilise metal ions to facilitate both their retention on the cell surface, as well as lattice assembly^24–31^. Thirdly and intriguingly, S-layer insertion is localised at the mid-cell and cell poles in many prokaryotes, including archaeal^32^, Gram-positive^33^ and Gram-negative bacterial species^4, 32, 34^.

One of the best understood systems for studying bacterial S-layers is provided by the model organism *Caulobacter crescentus*^35^. The S-layer of *C. crescentus* is comprised of a single SLP called RsaA^36^, which has the prototypical bipartite arrangement of SLPs^4^. We have reported the X-ray structure of the C-terminal domain of RsaA (RsaA_CTD_), consisting of residues 250- 1026, which form the highly-interconnected outer S-layer lattice^29^, and solved the cryo-EM structure of the N-terminal domain (RsaA_NTD_, residues 1-249) in complex with the O-antigen of lipopolysaccharide (LPS)^28, 37^, on which the S-layer is anchored^38^. S-layers of multiple species assemble in a metal-ion dependent manner^24, 26, 27, 30, 39^. Likewise, *C. crescentus* requires high concentration of extracellular calcium ions for SLP oligomerisation and retention on the cell surface^31^. Further, new S-layer insertion in *C. crescentus* is localised at the mid-cell and cell poles, by a mechanism that is not yet understood^31, 34^. In general, S-layer-expressing prokaryotes synthesise SLPs at incredibly high levels, and in the case of *C. crescentus*, RsaA has been suggested to account for between 10-31% of total protein content of the cell^40, 41^ and appears to be tightly regulated to prevent cytoplasmic build-up of excess protein^42^. Given the material and energetic demand imposed on the cell by S-layer production, it is reasonable to expect that S-layer assembly is carefully regulated.

In this study, we have investigated the cell cycle dependence of S-layer biogenesis in *C. crescentus*, using fluorescent microscopy and electron cryotomography (cryo-ET) of cells. Our results show that S-layer biogenesis is tightly linked with the cell cycle. We provide evidence showing that cell division and cell envelope biogenesis are regulated at multiple levels, providing new insight into the exciting field of S-layer biology, offering clues to why all S-layers (thus far) appear to be inserted at discrete locations in the cell.

## RESULTS

### S-layer insertion is localised to regions of cell-cycle dependent envelope growth in *C. crescentus*

To understand the cell-cycle dependency of S-layer insertion, we utilised a dual-labelling approach previously described in our study of calcium binding by RsaA^31, 43^. Briefly, to distinguish between old and newly inserted regions of the S-layer, we pulse-saturated the surface of *C. crescentus* cells expressing RsaA-467-SpyTag with SpyCatcher-mRFP1 (SC- mRFP1), followed by washing and chase labelling with SpyCatcher-sfGFP (SC-sfGFP) during exponential growth. Following labelling, we observed *C. crescentus* cells with distinct fluorescent regions of mRFP1- and sfGFP-labelling, corresponding to the pulse and chase respectively (Fig. 1A). Cell populations were asynchronous and fluorescence profiles varied depending on the cell size and cell cycle stage. Swarmer cells show limited or polar sfGFP labelling (Fig. 1B), while pre-divisional cells had a strong mid-cell sfGFP signal (Fig. 1C), in agreement with previous reports^31, 34^. As a control, *C. crescentus* cell cultures were briefly synchronised using density centrifugation with Percoll (Materials and Methods) and pulse-chase labelled as above, resulting in a similar labelling pattern to non-synchronised cells but with slightly larger regions of new S-layer (Supplementary Fig. 1).

**Fig. 1.**
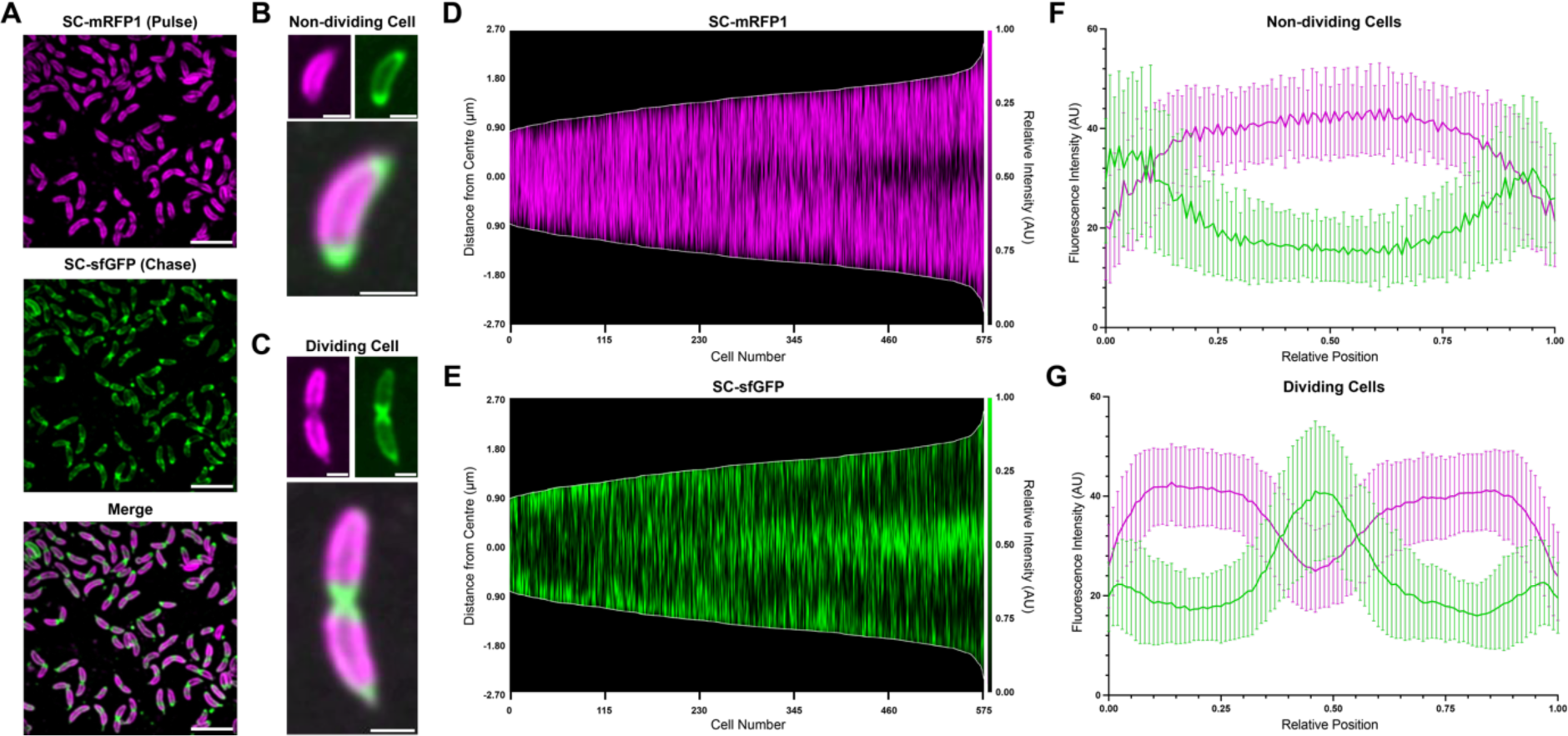
Incorporation of RsaA into the growing S-layer of *C. crescentus* relocates from the cell poles to the mid-cell during cell development. (**A**) Micrographs of *C. crescentus* RsaA-467-SpyTag cells pulse-chase labelled using SC- mRFP1 (top), SC-sfGFP (middle). Merged channels (bottom) show distinct localisation of the two SC-conjugates along the cell surface. Micrographs were gaussian filtered to remove noise. Scale bar = 5 µm. (**B-C**) Micrographs (SC-mRFP1 (top left), SC-sfGFP (top right), and merged channels (bottom)) of a representative non-dividing and dividing *C. crescentus* cell. Scale bar = 1 µm. Polar or mid-cell localisation of newly inserted S-layer being more prominent in non-dividing and dividing cell populations, respectively. (**D-E**) Demograph showing normalised fluorescent profiles of dual-labelled *C. crescentus* cells (n = 575), ordered by ascending length. (**D**) SC-mRFP1 signal corresponds to old S-layer, while (**E**) SC-sfGFP signal represents new S-layer. Shorter, swarmer cells show a propensity for polar localisation of new S-layer, while longer cells show mid-cell localisation of the sfGFP signal. (**F-G**) Relative intensity profiles of mRFP1 and sfGFP in (**F**) non-dividing cells and (**G**) dividing *C. crescentus* cells (n = 100 for both plots). Points were selected across the medial axis of each cell, and the normalised signal plotted by relative position along the cell. Error bars denote standard deviation.

To determine the relationship between S-layer localisation and cell size, cell profiles (culture shown in Fig. 1A) were ordered according to cell length in MicrobeJ using previously described methods^44^, and their fluorescent signals were then visually inspected (Figs. 1D-E). This analysis revealed a clear temporal progression of new S-layer formation (labelled with sfGFP signal) from poles to mid-cell. To further quantify the relationship between the cell cycle stage and the labelling pattern, the fluorescence profiles of non-dividing swarmer cells (cell length<2 µm) and dividing cells (assigned by the presence of a mid-cell invagination) were normalised and plotted against relative cell length. As expected from the visual inspection of the data, non-dividing cells (Fig. 1F) showed a stronger normalised sfGFP signal at their poles, while dividing cells had a much stronger signal at their mid-cell (Fig. 1G). The *C. crescentus* stalk, which is found on the pole of the dividing cell, was also encompassed by an S-layer^29, 36^. Stalk biogenesis represents a rapid transition from swarmer to sessile cell type in *C. crescentus*, the latter representing the dividing population^45–47^. Labelling patterns of the cell stalks appeared to be dependent on whether stalks were present during the pulse or were generated during the chase-labelling. Although this led to variable and occasionally dual-coloured stalks, even when cells were synchronised, the base of the stalk near the cell body always labelled positive for new S-layer insertion (Supplementary Fig. 1).

### Inhibition of cell-division does not cause delocalisation of S-layer insertion

To further explore what underlying processes underpin the observed localisation of RsaA insertion into the growing S-layer, we sought to perturb the cell cycle of *C. crescentus* by using compounds that either inhibit cell division or cell elongation, to assess the local accumulation of new and old S-layer material. The distinctive regions of old (mRFP1) and new (sfGFP) S-layer observed using our pulse-chase method (Fig. 1), were quantified for co-localisation using Pearson’s Correlation Coefficient (PCC)^48, 49^. PCC quantifies the linear correlation between two datasets (in this case, two channels of a fluorescence image): a PCC > 0 suggests there is colocalisation between the two channels, whereas PCC < 0 shows anticorrelation^48, 50–52^. As expected from visual inspection of our data (Fig. 1), in the absence of cell-cycle perturbing compounds, pulse-chase labelled *C. crescentus* cells exhibited a strong anticorrelation (average PCC = -0.46) between the old and the new S-layer regions (Figs. 2A and 2D), consistent with our previous observations (Fig. 1).

**Fig. 2.**
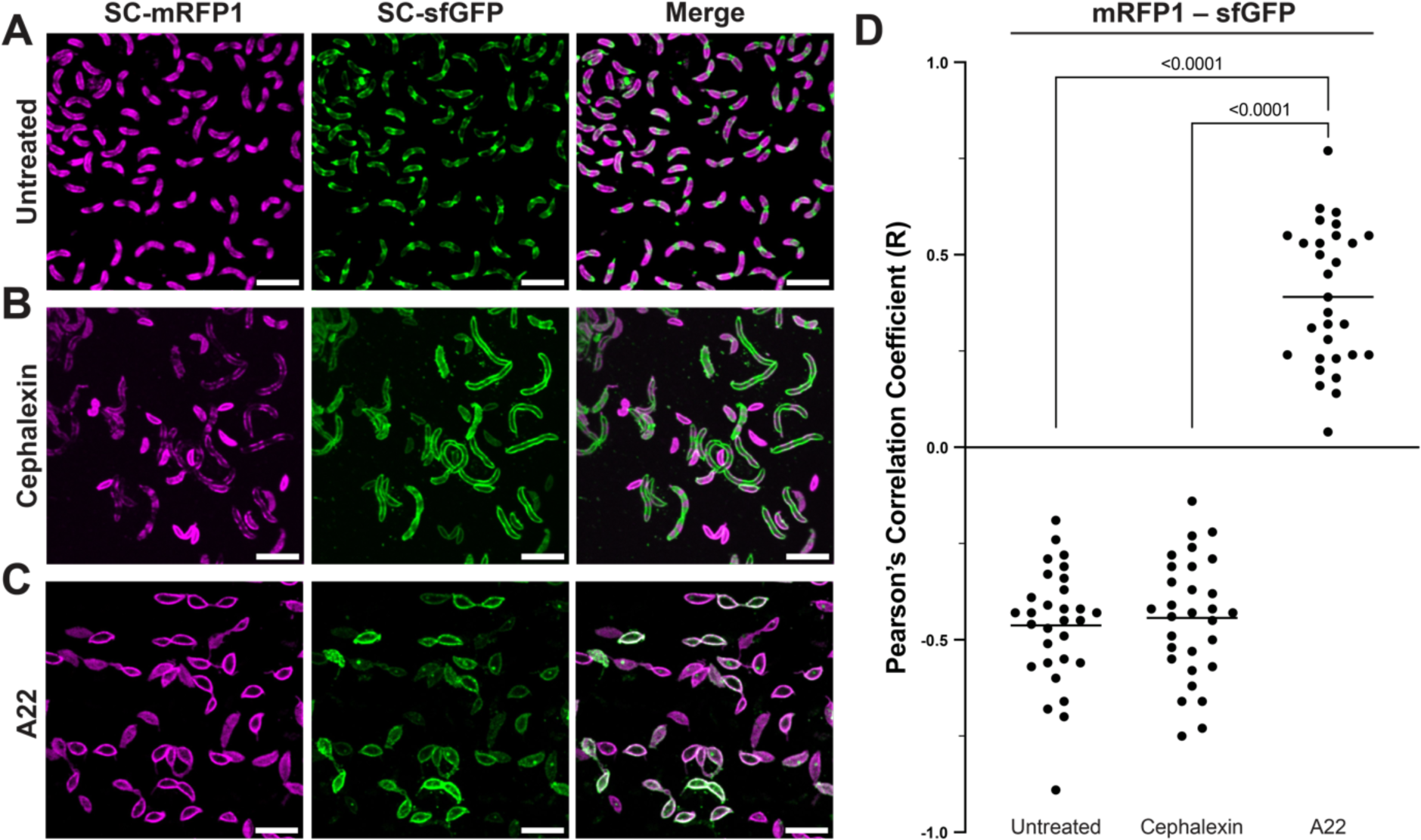
Inhibition of MreB, but not cell division, results in delocalisation of S-layer proliferation in *C. crescentus*. (**A-C**) Micrographs of dual-labelled *C. crescentus* RsaA-467-SpyTag cells. SC-mRFP1 (left), SC-sfGFP (centre) and merged channels (right). Scale bars = 5 µm. Cells were pulse-chase labelled using the same procedure to that of Fig. 1, but under varying conditions, including (A) no treatment, (B) cells treated with 50 µg/mL cephalexin, and (C) 3 µg/mL A22. (D) Analysis of colocalisation by PCC show untreated and cephalexin treated cells have a negative colocalisation (R value) between the mRFP1 and sfGFP channels, as expected given the visually observable separation of old and new S-layer. A22 treated cells show a positive R value and colocalisation between the both channels, suggesting MreB inhibition has resulted in the loss of discrete localisation of RsaA insertion into the S-layer during cell growth. One-way ANOVA analysis of the data shows a strong significant difference between A22 treated cells and both the other conditions (Student’s *t*-test, p <0.0001), whereas comparing untreated and cephalexin treated cells showed no significant differences. N = 30 cells analysed for all treatment conditions.

To test whether local S-layer assembly depends on the cell division machinery, we next treated cells with cephalexin, a cephalosporin antibiotic that inhibits cell division by disrupting peptidoglycan (PG) synthesis at the mid-cell during division^53–55^. Exposure to sublethal concentrations of cephalexin inhibited cell division, and resulted in the formation of lines of connected cells in filaments (Fig. 2B), consistent with published work^56^. These filamentous cells were labelled in the same manner as untreated cells, but with a prolonged chase (3 hours) to allow for growth. Despite cephalexin’s profound impact on cell viability, cephalexin-treated cells retained a dual-labelled S-layer pattern with distinct regions of old and new S-layer (Figs. 2B and Supplementary Fig. 2). Repeating the co-localisation analysis in these cells confirmed that the old and new S-layer regions were strongly anticorrelated, almost to the same extent as untreated cells (Fig. 2D, average PCC = -0.45, no significant difference in Student’s *t*-test). To confirm whether the S-layer in these drug-treated cells had a normal appearance, we vitrified the cephalexin treated cells on electron microscopy (EM) grids and imaged them using cryo-ET. Reconstructed tomograms confirmed that the S-layer is positioned ∼18 nm away from the outer membrane, forming a hexagonal lattice (Figs. 3A-B and Supplementary Fig. 3), consistent with tomograms of untreated cells in published data^31^. These results together suggest that cephalexin treatment does not affect S-layer secretion or localised assembly.

**Fig. 3.**
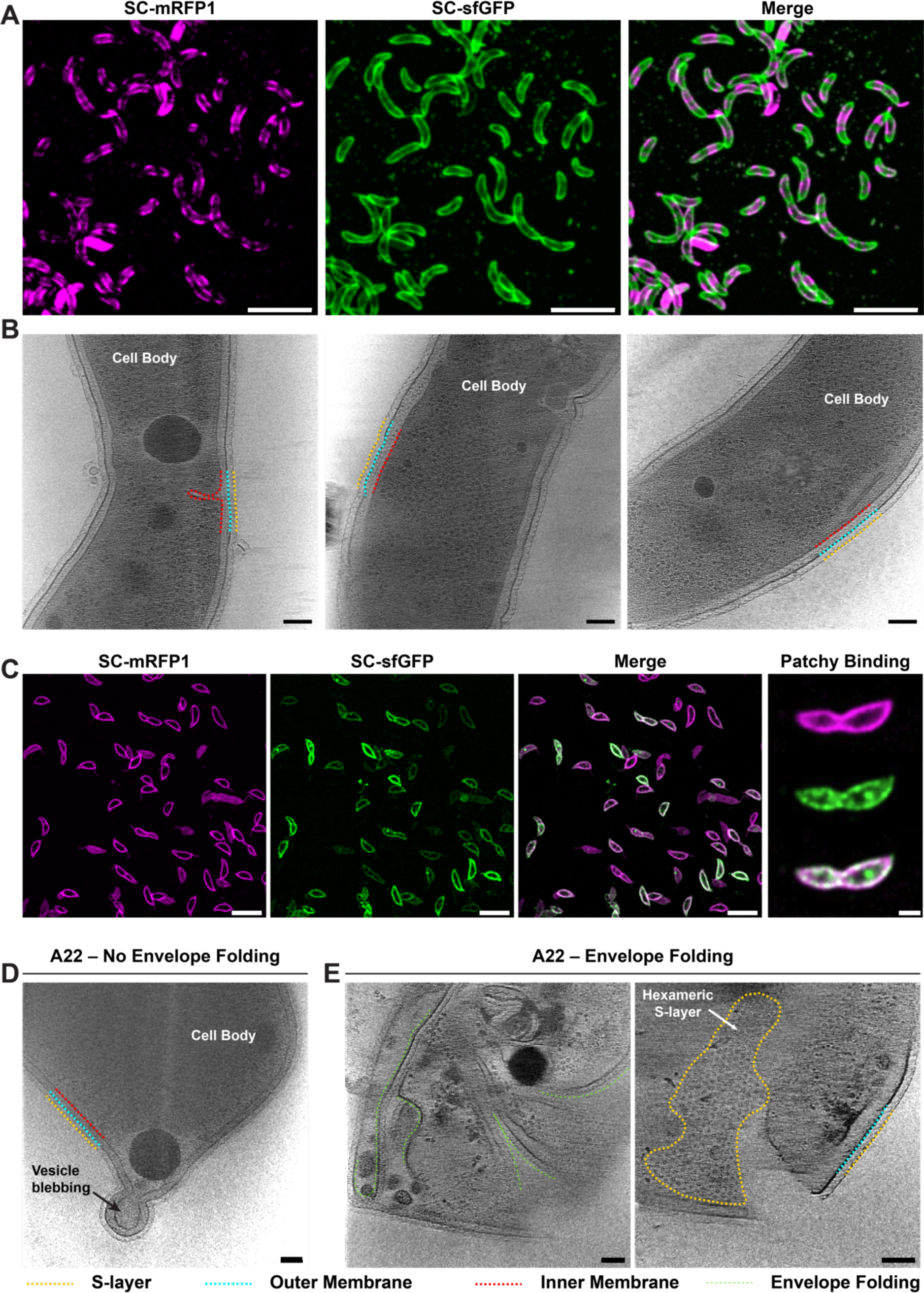
Drug-treated cells retain a continuous S-layer bound to the OM. (**A**) Micrographs of dual-labelled, cephalexin-treated *C. crescentus* RsaA-467-SpyTag cells, SC-mRFP1 (left, magenta), SC-sfGFP (centre, green) and merged channels (right). Scale bar = 5 µm. (B) Slices through tomograms of vitrified cephalexin-treated *C. crescentus* cells, showing a complete S-layer bound to the OM. components of the cell envelope are labelled (Yellow-S-layer, cyan-OM, red-IM)). Scale bar = 50 nm. (**C**) Micrographs of dual-labelled, A22 treated *C. crescentus* RsaA-467-SpyTag cells, same labels as panel A. Scale bar = 5 µm. On the extreme right is an example of an A22-treated cell showing “patchy” S-layer labelling with overlapping sfGFP and mRFP1 signals. Scale bar = 1 µm. (**D-E**) Slices through tomograms of vitrified, A22-treated *C. crescentus* cells. (D) Enlarged cell showing budding vesicle forming at the presumed cell pole. (D) Deformed cells showing folding of envelope as a result of A22 exposure. Folds in the envelope have been labelled (green). The second panel shows a higher Z-slice of the same reconstructed tomogram, where the top view of the S-layer is clearly visible and adopts a regular hexagonal arrangement. Scale bar = 50 nm. Components of the cell envelope have been labelled as in panel **B**.

### Disruption of the cytoskeletal protein MreB delocalises S-layer insertion

Having shown that cell division has little impact on the localisation of S-layer biogenesis, we next investigated the effect of disrupting cell elongation and rod morphogenesis by the disruption of the bacterial actin homologue MreB. To do so, we treated *C. crescentus* cells with the compound A22, which has been shown to bind to MreB and disrupt cell shape^57, 58^ and cell polarity^59, 60^. We then investigated the effect of disruption of MreB filaments on S-layer biogenesis. Strikingly, S-layer integration at the surface of these A22-treated cells was delocalised (Fig. 2C-D). Furthermore, pulse-chased labelled cells adopted a “lemon”-shape and showed several regions of new S-layer without the previously observed mid-cell or polar localisation. Quantification of this co-localisation (Fig. 2D) confirmed that this treatment led to loss of the anticorrelation between new and old S-layer (average PCC = 0.44). Furthermore, the patterns of new and old S-layer localisation observed in these experiments were significantly different from those seen in control untreated and cephalexin-treated cells (p <0.0001 in pairwise Student’s *t*-tests).

Broadly, two different types of labelling patterns were observed in “lemon”-shaped A22- treated cells (Fig. 3C and Supplementary Fig. 2). The first type exhibited almost no new S-layer labelling, so that cells only stained for old S-layer, and a second type in which old and new S-layers both appeared delocalised (Fig. 2C and 3C). To better understand these labelling patterns and to examine ultrastructure of the S-layer in the A22-treated cells, we performed cryo-ET of treated cells that were vitrified on EM grids (Fig. 3D-E). Tomograms confirmed the “lemon”-shaped appearance of the cells, and the presence of an S-layer on the surface with similar morphological parameters as the wild-type S-layer (Fig. 3D-E). In line with the fluorescent microscopy, tomograms also confirmed two phenotypes, both of which were “lemon”-shaped and possessed an S-layer with similar morphological parameters to that of the wild-type S-layer (Fig. 3D-E). One set had severe cellular disruption including invaginated membranes (Supplementary Fig. 3).

### Cell wall turnover precedes S-layer biogenesis at the mid-points of dividing cells

Given the known dependence of PG biogenesis on MreB in *C. crescentus*^61^, we next explored the relation between new PG and new S-layer by labelling newly synthesized PG using fluorescent D-amino acids^62^ alongside old and new S-layer. PG labelling with HADA (a blue fluorescent D-amino acid) showed distinct fluorescent punctae in cells (Fig. 4A), seen previously in several bacteria^62^, including *C. crescentus*^63^. A visual inspection of fluorescent images suggested the co-localisation of new PG and new S-layer insertion. To test this hypothesis, we obtained profiles along the length of each cell and ordered each cell according to their lengths (Fig. 4B-D). This analysis revealed that HADA fluorescence was localised at the mid-cell in short cells in earlier stages of the cell cycle, while the integration of new S-layer material, as indicated by the presence of sfGFP, occurs in longer cells at later stages of the cell cycle, suggesting PG turnover precedes S-layer biogenesis (Fig. 4B-D and Supplementary Fig. 4). New S-layer insertion begins at the mid-cell in dividing cells, regions where no old S-layer is detected. In the longest cells analysed, new PG insertion was not detected, despite the presence of new S-layer insertion, indicating that PG insertion concludes around the time new S-layer insertion begins.

**Fig. 4.**
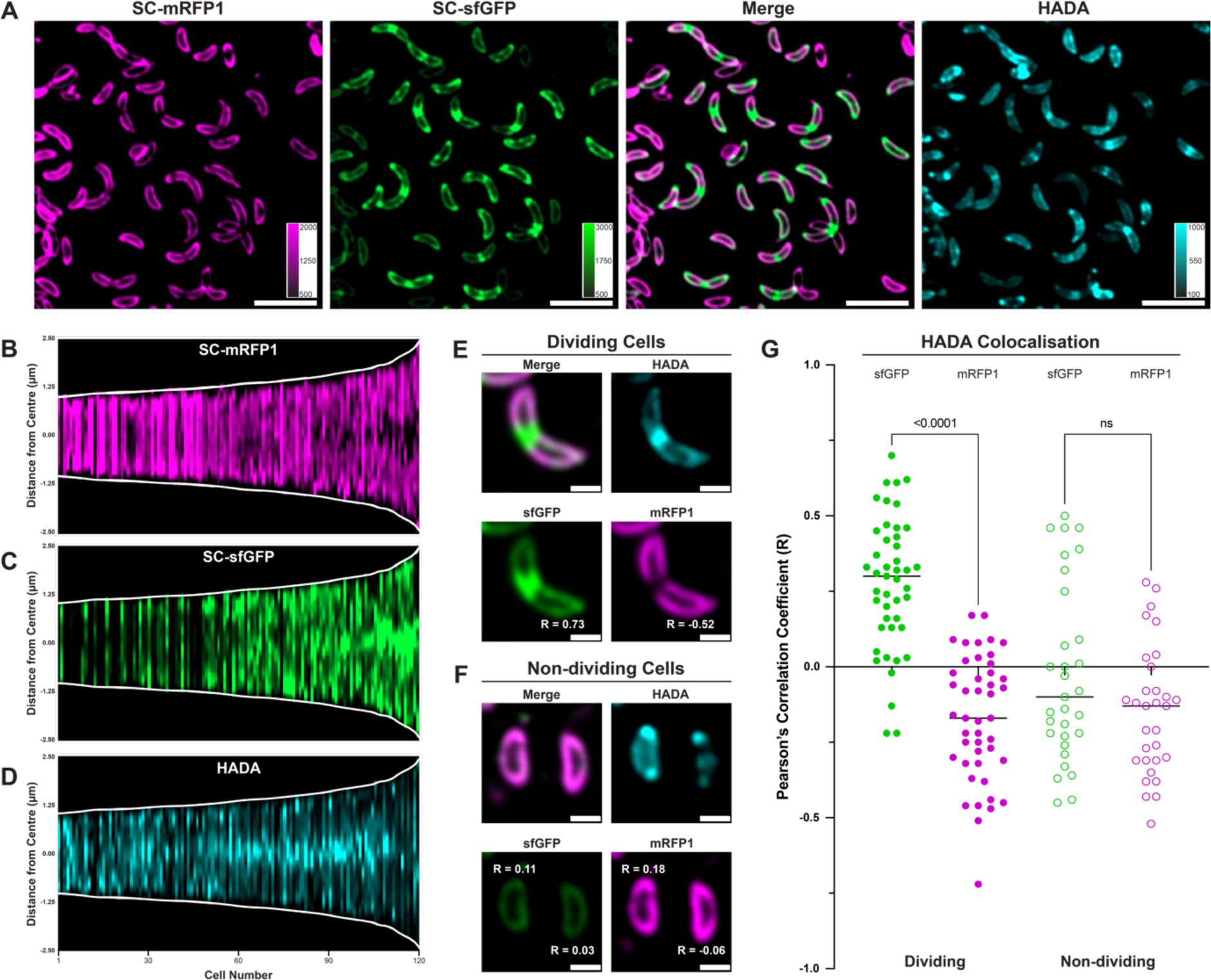
Cell wall turnover precedes and colocalises with S-layer expansion in dividing *C. crescentus* cells. (**A**) Three-colour labelled *C. crescentus* cells, pulse-labelled with SC-mRFP1 (magenta), chased with SC-sfGFP (green). Channels showing S-layer labelling are merged (third panel) and the HADA fluorescence shown in cyan (fourth panel). Calibration bars are provided for each channel. Scale bars = 5 µm. (**B-D**) Demographs of labelled *C. crescentus* cells ordered by length (n = 100), showing (**B**) SC-sfGFP, (**C**) SC-mRFP1, and (**D**) HADA fluorescence profiles for each cell. HADA signal localises to the mid-cell at shorter cell lengths than SC-sfGFP, which precedes an apparent colocalization of the sfGFP and HADA signals. The HADA signal at mid-cell eventually subsides, prior to further new S-layer insertion. Demograph intensities are calibrated to the same levels as in A. (**E**-**F**) PCC scores (R) for correlation of HADA with sfGFP and HADA with mRFP1 signals, comparing (**E**) dividing and (**F**) non-dividing cells. R values are displayed next to their respective cells in the relevant channels. Scale bars = 1 µm. (**G**) PCC scores (R) between HADA and sfGFP or mRFP1 channels for dividing (n = 45) and non-dividing (n = 31) cells measuring colocalisation. Dividing cells show a significantly higher R value between HADA and sfGFP (_●_) compared to mRFP1 (_●_) (measured by Student’s *t*- test), suggesting stronger colocalisation. Non-dividing cells showed a negative PCC R score on average for HADA correlation between both sfGFP (**_○_**) and mRFP1 (_○_) signals, with no significant difference between the two.

To confirm quantitatively that new PG and new S-layer is inserted in the same locations in cells, we repeated the co-localisation analysis described above (Fig. 2), measuring HADA fluorescence co-localisation with both new and old labelled S-layer (Fig. 4E-G). In non-dividing cells, there was no significant difference between the co-localisation measured between new PG and new or old S-layer (PCC = -0.10 new PG / new S-layer and PCC = -0.12 new PG / old S-layer). In contrast in dividing cells, new PG was co-localised with new S-layer (PCC = 0.30) rather than with old S-layer (PCC = -0.15), in a statistically significant difference (Student’s *t*-test, p-value < 0.0001). These observations suggest that cell wall expansion is a driving force in cell envelope growth and a predictor of local S-layer biogenesis.

### S-layer insertion occurs at gaps in the S-layer lattice

Having studied the cell-cycle dependence of S-layer biogenesis, we next scrutinized how new RsaA molecules insert themselves into a pre-existing two-dimensional lattice packed with proteins spanning the cell envelope. For this, we utilised cryo-ET data of *C. crescentus* cells, focusing on the mid-cell, as done previously^64, 65^, or the cell poles (Fig. 5), i.e. regions of the cell where we have shown the new S-layer is inserted (Fig. 1). Cryo-ET allowed us to observe the ultrastructure of the S-layer, allowing us to go beyond the diffraction-limited optical microscopy pictures to study the morphology of the new S-layer insertion sites (Fig 5). Unexpectedly, we observed gaps in the S-layer lattice at the S-layer biogenesis sites. These were seen in all tomograms analysed (Fig. 5A-F). As a control, we vitrified dual S-layer labelled *C. crescentus* cells on EM grids (see Fig. 1) for cryo-correlated light and electron microscopy (cryo-CLEM). Cryo-light microscopy of the vitrified cells, although limited in resolution in our widefield setup, allowed us to identify cells with clear dual labelling and new S-layer insertion (Supplementary Fig. 5A-B). These cells were then located in the electron microscope by overlaying the light microscopy images with overview images of EM grid squares. Cryo-ET of these dual labelled cells confirmed gaps in the S-layer at the site of new S-layer insertion (Supplementary Fig. 5C-E), confirming our cryo-ET observations (Figs 5A-F). These gaps are locations of discontinuity of the S-layer, where either two lattices appear to overlap or rows of hexamers are missing (Fig. 5A-F). At the several areas, two separate two- dimensional sheets of S-layers appear to meet, showing up as a line defect on the cell surface. Given their placement at regions of cell-envelope expansion, these likely represent regions of RsaA insertion or lattice formation.

**Fig. 5.**
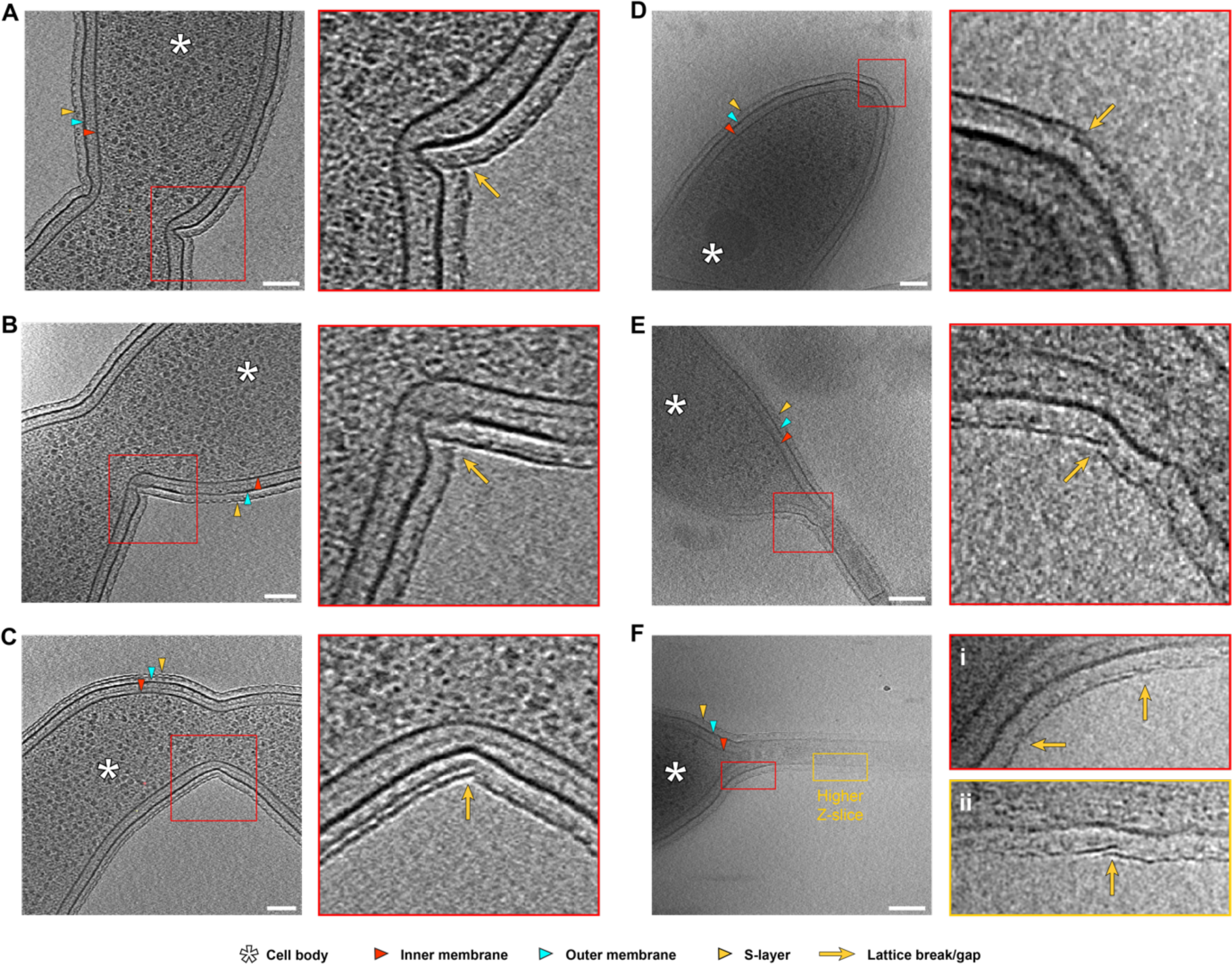
S-layer insertion events at regions of cell growth and high membrane curvature identified in our light microscopy experiments above. (**A-F**) Slices through tomograms of *C. crescentus* cells (left panel) and zooms defined by the red square (right panel) showing possible insertion events. Insertion events cover regions of S-layer biogenesis, as established by our light microscopy. Tomograms in **A-C** show mid-cell discontinuities of the S-layer, **D** shows the flagellate cell pole (the flagellum is visible at lower Z-slices), **E-F** show *C. crescentus* stalks. Components of the cell enveloped have been labelled – a key for the symbols used to identify the cell body, IM, OM, S-layer and insertion/overlapping regions is provided at the bottom of the figure. Specifically, yellow arrows denote regions of discontinuities or overlaps in the S-layer lattice. Cases where the Z-slice have been changed for the zoomed panel have been marked. Scale bars = 100 nm.

## DISCUSSION

Based on our analyses, we suggest a new model of S-layer biogenesis dependent on the cell cycle (Fig. 6). We suggest that areas of cell growth that contain new membranes and potentially freshly secreted LPS molecules, which do not assemble precoated with RsaA, likely resulting in gaps in the S-layer (Fig. 6). Additionally, regions of cell growth in *C. crescentus* often contains significant membrane curvature, which likely contribute to shear stress in the S-layer leading to lattice rupture, because geometrically, a hexagonal lattice cannot tesselate perfectly along curved surfaces. A pool of RsaA molecules is known to be present between the OM and the S-layer, freely diffusing on the LPS, evidenced by light microscopy^34^, cryo-EM structures and cryo-ET of cells^28^. These unassembled RsaA molecules would always be available to plug any gaps in the lattice, caused by damage from environmental pressures or, as observed in our study, regions of cell growth and high membrane curvature. This allows the cell to retain a nearly complete S-layer as it moves through the cell cycle.

**Fig. 6.**
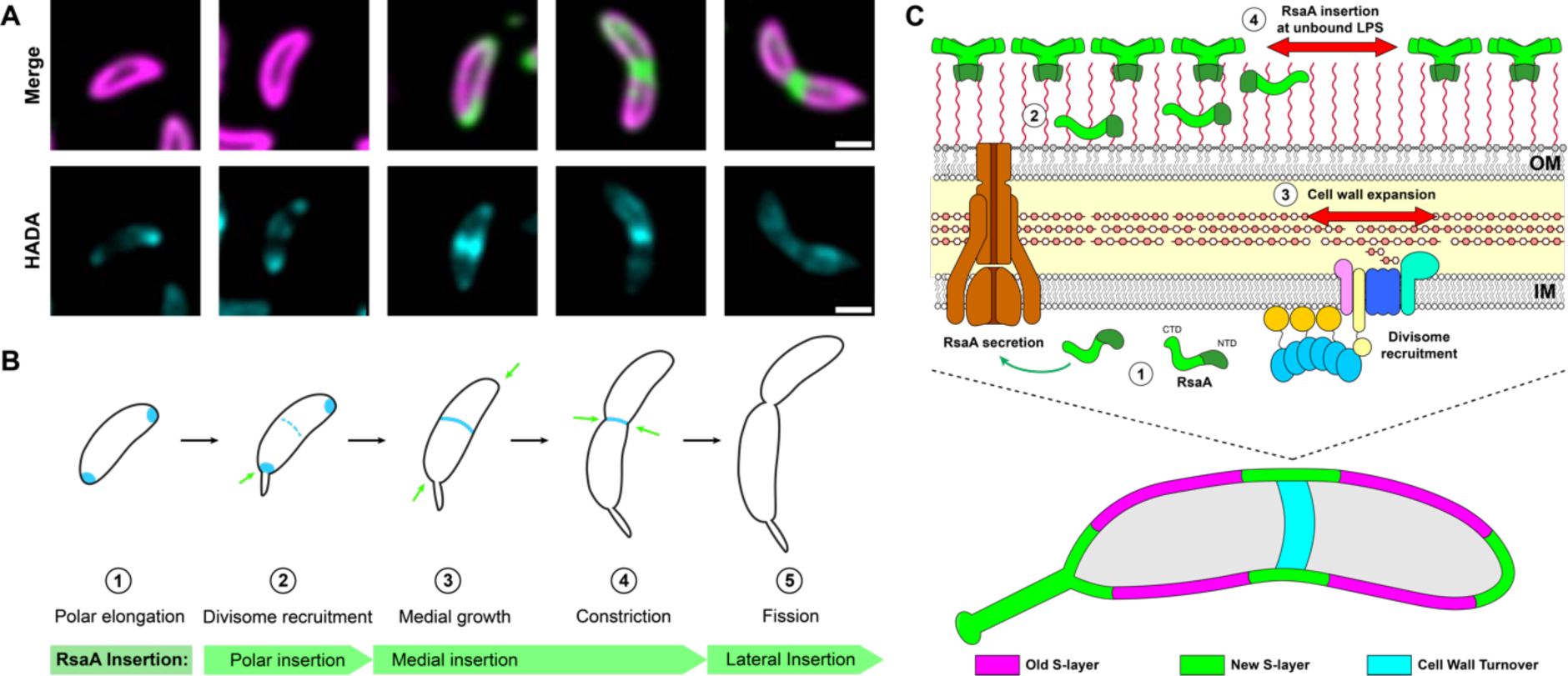
Proposed model of cell-expansion dependent S-layer insertion in *C. crescentus*. (**A**) Micrographs showing individual *C. crescentus* cells of varying length, beginning with a swarmer cell and ending with pre-divisional cell. Top row shows merged SC-mRFP1 and SC- sfGFP signals, bottom row shows HADA signal. Scale bars = 1 µm. (**B**) Schematic representation of the relationship between re-localisation of S-layer biogenesis to regions of cell wall turnover and expansion. (**C**) Model of S-layer biogenesis in *C. crescentus*; unfolded, monomeric RsaA is exported from the cytoplasm into the extracellular space, whereupon it binds LPS. RsaA then diffuses along the LPS until it finds a region of surface expansion corresponding to a gap in the S-layer, driven by PG turnover and polymerisation in the cell wall, leading to RsaA binding to the tip of the LPS and oligomerising with the pre-existing S- layer to complete the lattice.

The localisation pattern of S-layer biogenesis as observed by our SpyCatcher-labelling approach is remarkably similar to the localisation patterns of several proteins key to cell division. For example, fluorescently labelled components of the divisome machinery in *C. crescentus*, such as DipM (LytM endopeptidase), FtsW (PG polymerase), FtsL (divisome- recruitment protein), and PBP3 (PG-crosslinking divisome protein), share marked similarities with new S-layer biogenesis^55, 66–68^. Unlike S-layer biogenesis, the recruitment of these components is much better understood and relies on the highly-conserved prokaryotic tubulin homologue, FtsZ, which, along with several other key proteins, comprises the divisome complex in *C. crescentus*^64, 67, 69–71^. As successful cytokinesis requires the breaking and remodelling of the PG cell wall, many of these division proteins are associated with significant cell wall turnover, and would therefore likely co-localise with new S-layer insertion^56, 72–74^.

These past studies on cell division and our results here suggest a multi-level co-ordination for the homeostasis of the cell envelope within the *C. crescentus* cell cycle (Fig. 6). There is clear tendency for bacterial and archaeal cells to synchronise the biogenesis different envelope components^4^, which appears to also be the case in *C. crescentus* (Fig. 6). While it is tempting to suggest that unbound LPS in the observed gaps is freshly secreted, remarkably little is known regarding the potential localisation of LPS integration into the OM in *C. crescentus*. In two Gram-negative bacteria *Brucella abortus* and *Agrobacterium tumefaciens* that exhibit polar growth^75^, localisation of the LPS-biosynthetic machinery to regions of cell-growth has been demonstrated^76^. Additionally, previous studies have shown that the OM is diffusion restricted and the OM composition is directly regulated by cell wall turnover in bacteria such as *Escherichia coli*^77^. Polymerised *C. crescentus* S-layer (but not monomeric RsaA), also appears to be diffusion-restricted, so it is possible that these SLP-deficient OM may also have been recently inserted in *C. crescentus*.

Owing to the crystalline nature of the *C. crescentus* S-layer, SLPs are likely able to self- integrate themselves into the growing lattice at gaps in the two-dimensional crystal. This has been observed *in vitro*, where purified RsaA in the presence of calcium spontaneously forms hexameric lattices comparable to those seen on the cell surface^29, 30, 78^. This presents an ingenious solution to a difficult logistical problem of the polymerisation of the highly ordered S-layer, as it requires no further energetic input from the cell beyond secretion of the constituent SLP into the extracellular milieu, where it will bind the surface and oligomerise. It is remarkable therefore, that other studied S-layers with a similar mid-cell insertion phenotype, do not possess extensive gaps in the lattice, but rather complete the S-layer with defined geometric defects^79–81^. For example, in the archaeal S-layer of *Haloferax volcanii*, pentamers and heptamers (but no gaps) were observed on cells, which were coated to near-perfect continuity by the hexagonal S-layer^12^. Geometrically, to close a hexagonal sheet, defects or gaps must be present, therefore it will be intriguing to study why different organisms have adopted different solutions to this problem. Future research in this direction will illuminate our understanding of curved lattices in cells, which are ubiquitous across domains of life.

Our study did not localise the RsaA secretion machinery, the type 1 secretion system (T1SS) RsaDEF, the components of which share homology with a variety of other T1SS machineries^38, 40, 41, 82–85^. While principles of egress by T1SS have been extensively investigated, there is little literature on the localisation of T1SS across Gram-negative bacteria^86–88^. Studies on the localisation of RsaDEF would provide further context into the mechanisms of S-layer insertion in *C. crescentus* and other species.

S-layers are widespread in prokaryotes, and fundamental biology related to S-layers is poorly understood. This cell biology study of S-layer insertion in *C. crescentus* attempts to address this important gap in our knowledge and will help place future studies on S-layers into context. Our results into S-layer biogenesis will also be of great interest to microbiologists studying cell division and the cell cycle because S-layer biogenesis appears to be tightly linked to the cell cycle in many organisms across domains of life^4^. Our studies also have implications in the design and utility of *C. crescentus* and RsaA S-layers as a platform for synthetic biology applications. Indeed, using our S-layer structural data, several applications have already been reported^43, 89^. These implications on fundamental microbiology and synthetic biology highlight why urgent future research is needed to understand these captivating two-dimensional arrays found in abundance in prokaryotes.

## SUPPLEMENTARY MATERIAL

**Supplementary Table 1.**
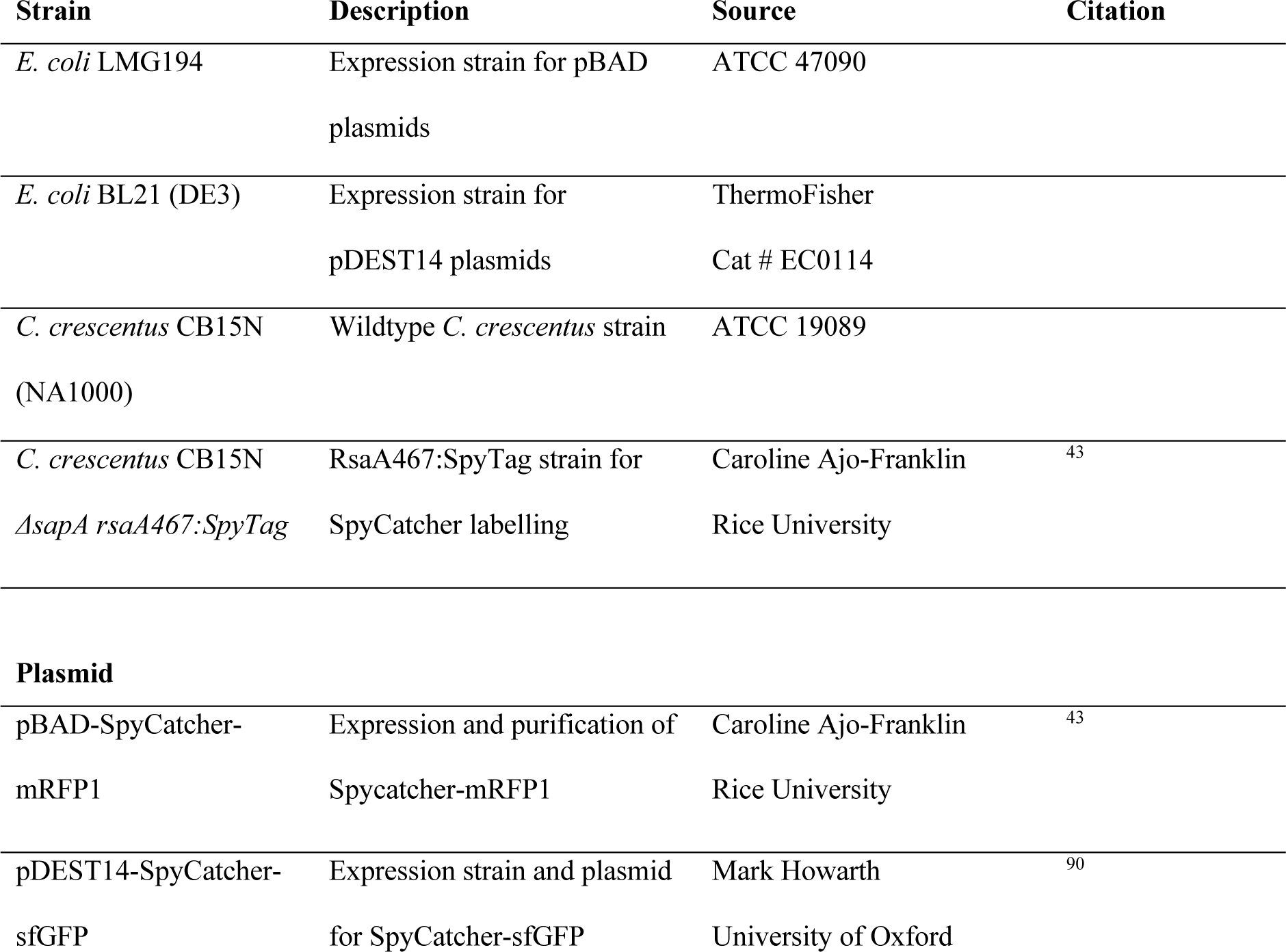
Strains and plasmids used in this study.

**Supplementary Fig. 1.**
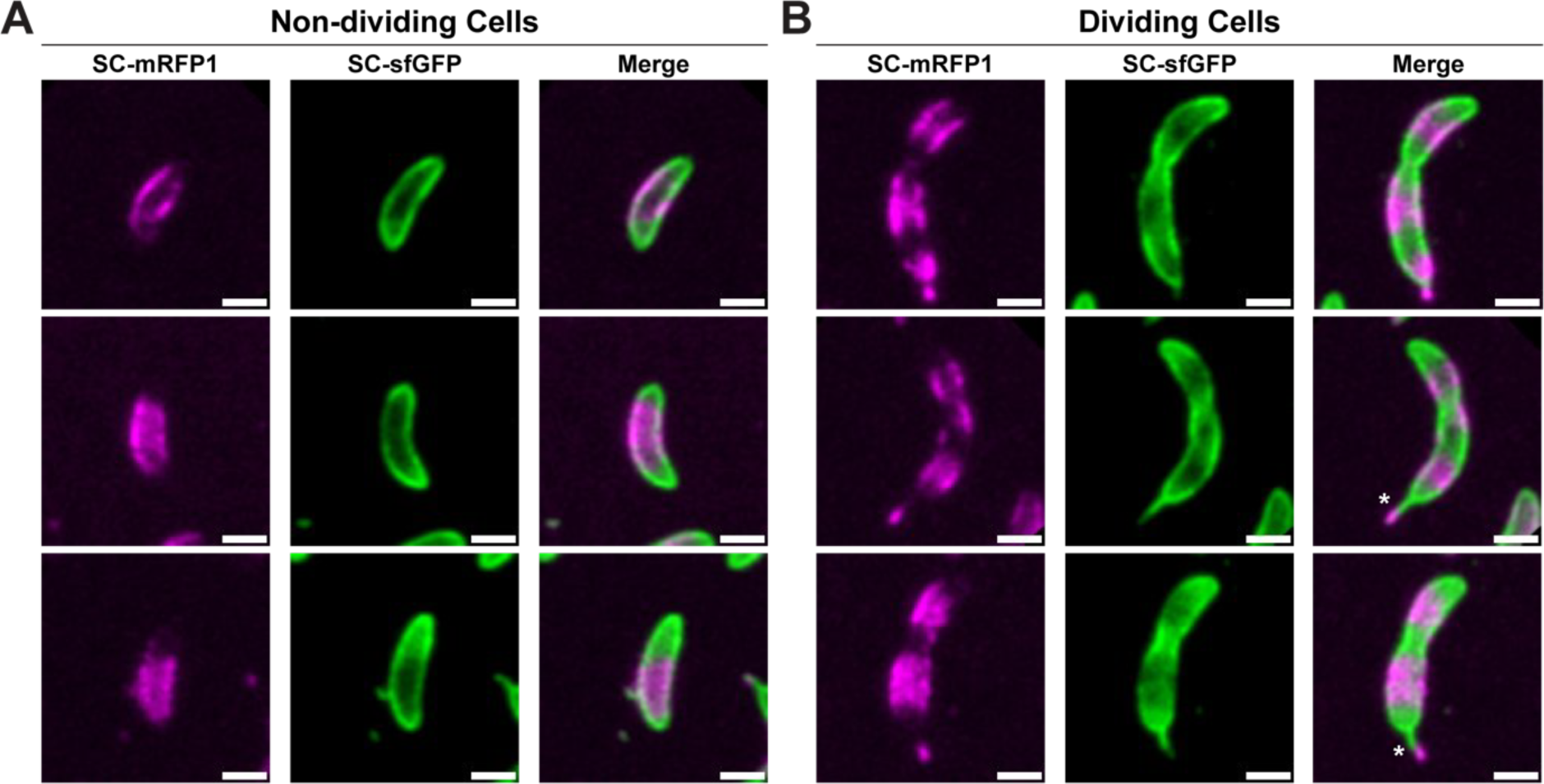
S-layer localisation patterns in dividing and non-dividing *C. crescentus* cells. Comparison of labelling in (**A**) non-dividing cells and (**B**) dividing, stalked cells. Cells were synchronised prior to pulse-chase labelling using SC-mRFP1 and SC-sfGFP as described. Polar labelling can be seen in all cells, but mid-cell labelling is only apparent in dividing cells. Additionally, dual-coloured stalks (SC-sfGFP at the base of the stalk, and SC-mRFP1 at the stalk tip) are indicated by an asterisk. This is consistent with previous research that shows stalk- biogenesis pre-empts cell division and new stalk material is created from the base^91^. Scale bars = 1 µm.

**Supplementary Fig. 2.**
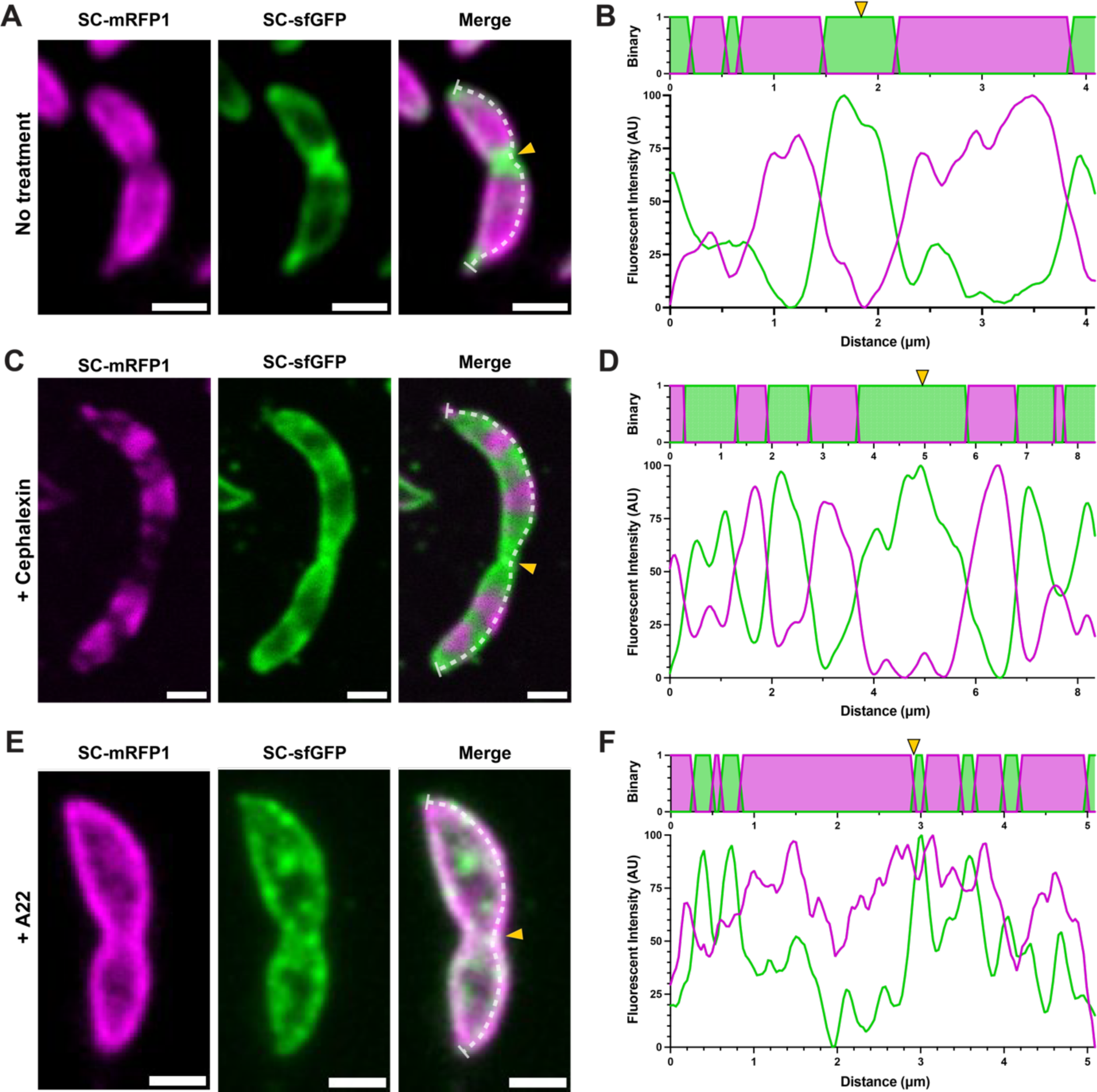
Cephalexin treated *C. crescentus* cells display a similar labelling pattern to untreated cells, markedly different from cells treated with A22. (**A**) mRFP1 (magenta), sfGFP (green) and merge micrographs of a representative untreated cell. (**B**) A three-pixel line was manually drawn along the right-axes of the cell (white dashed line) and the resulting profiles are displayed to the right of the micrographs. Scale bars = 1 µm. The fluorescence profile (starting from the northmost point of the cell) was normalised and plotted accordioning to the position along the cell axis. Above the normalised fluorescence data, a binary projection of the two channels is presented to show the dominant signal along the cell. The mid-cell, as determined by the presence of invagination, is indicated by a yellow arrow on the merged image and the binary cell profile. (**C-D**) Corresponding mRFP1, sfGFP and merge micrographs of a representative cephalexin-treated cell (50 µg/mL) along with the computed profiles. (**E-F**) Corresponding mRFP1, sfGFP and merge micrographs of a representative A22-treated cell (3 µg/mL) along with the computed profiles.

**Supplementary Fig. 3.**
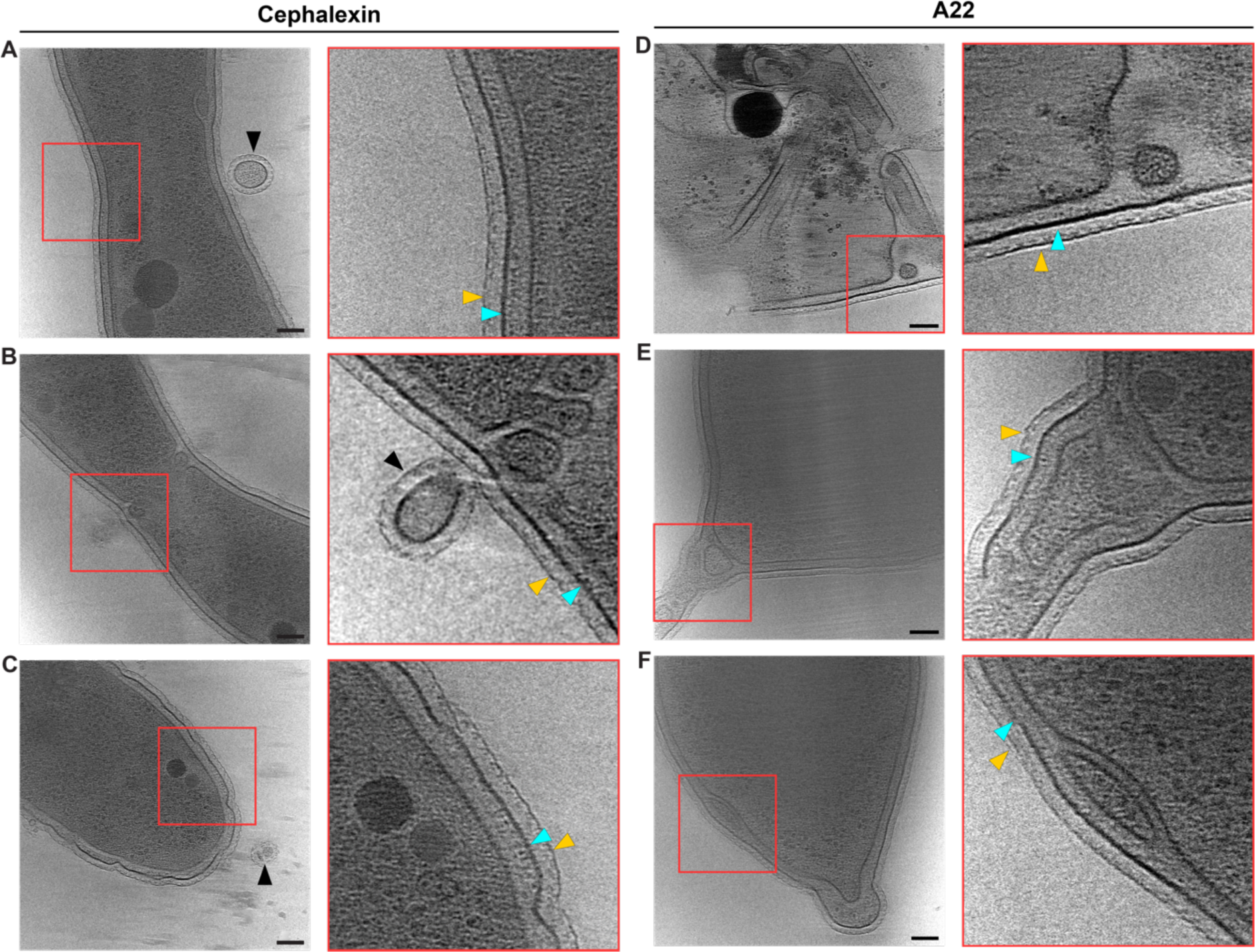
Cryo-ET gallery of cephalexin and A22 treated *C. crescentus* cells. Slices through reconstructed tomograms of (**A-C**) cephalexin (50 µg/mL) and (**D-F**) A22 (3 µg/mL) treated cells. The left panel in each case shows a view of the cell body (scale bars = 100 nm),while the right panel shows a zoomed-in view of the regions highlighted by the red box. Z-slice has been adjusted in the zoomed views to highlight the cell envelope. Yellow arrows indicate the S-layer, cyan arrows the OM.

**Supplementary Fig. 4.**
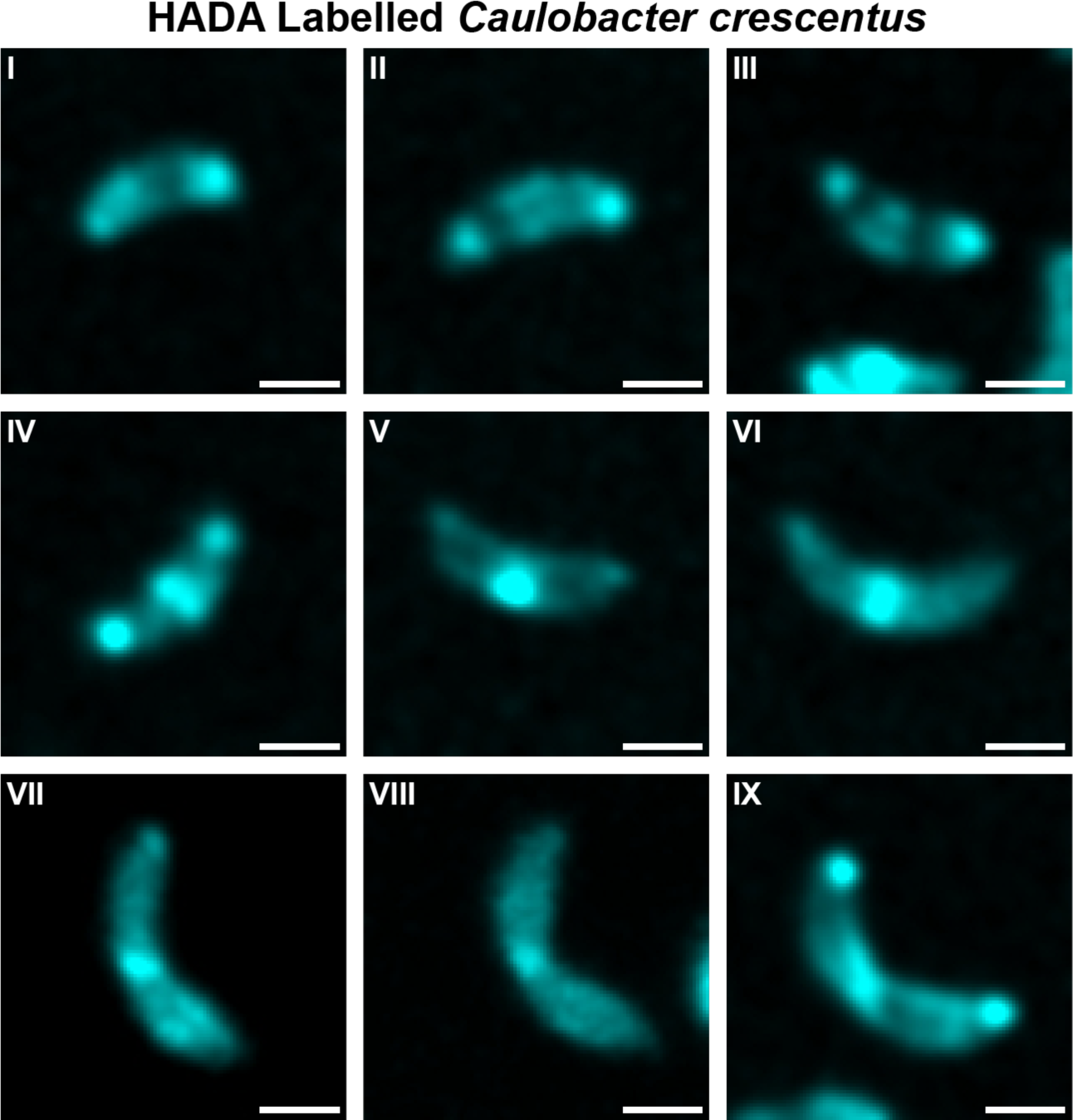
Gallery of HADA labelled *C. crescentus* cells (I-IX) Micrographs showing HADA-labelled *C. crescentus* cells arranged by ascending cell length. Scale bars = 1 µm.

**Supplementary Fig. 5.**
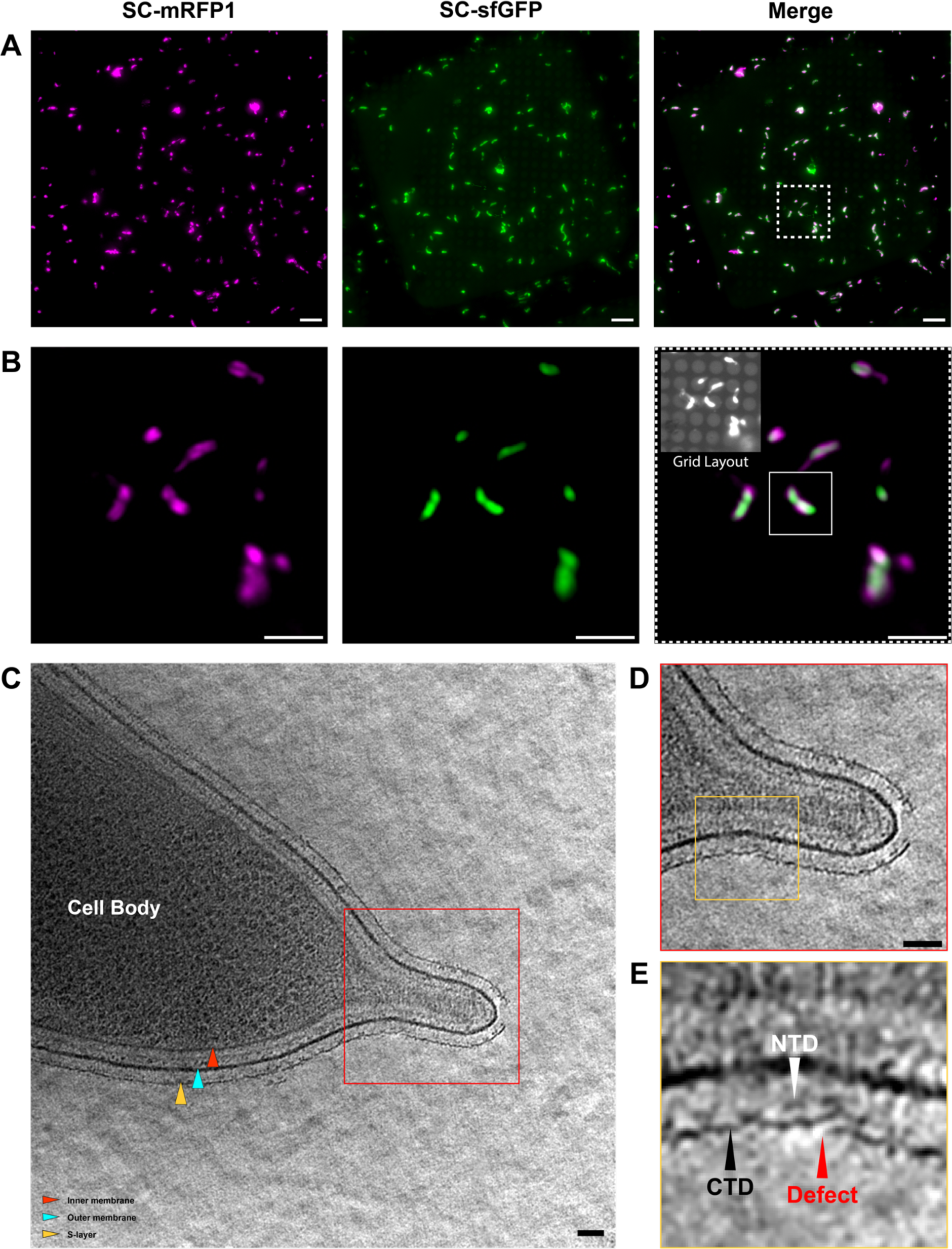
CryoCLEM experiment to confirm that RsaA inserts at gaps in the cellular S-layer. (**A**) Maximum Z-projection of a Z-stack through a cryo-EM grid containing vitrified, dual-labelled *C. crescentus* cells (labelled as in Fig. 1, scale bars = 10 µm). Regions of old S-layer are highlighted in magenta (SC-mRFP1), new S-layer in green (SC-sfGFP), and a merge of both channels is provided in the last panel. (**B**) A zoomed view of the subsection of the micrograph showing the region used for cryo-ET data collection, highlighted in the merged panel of A (white dashed border), channels are arranged as in A (scale bars = 5 µm). The inset in the merged channel micrograph shows a re-contrasted image where the layout of the EM grid, and the cell selected for cryo-ET collection is highlighted by a white box. **(C)** Slice through a tomogram of the dual-labelled *C. crescentus* cell highlighted in B. Components of the cell envelope are labelled using coloured arrows (legend in the bottom left of the panel). (**D**) A zoomed view of the region highlighted by the red box in **C**, showing a short stalk. Gaps in the S-layer can be seen at the stalk tip (likely due to the presence of the holdfast- polysaccharide^92^) and at the base of the stalk. Scale bars = 50 nm. (**E**) A further closeup of the latter is given in the bottom right panel, showing overlapping regions of the S-layer (labelled as “Defect” using a red arrow). The N-terminal and C-terminal domains (NTD and CTD) of RsaA in the assembled S-layer are marked by white and black arrows, respectively.

## Supplementary Movies

**Movie S1:** Tomogram of cephalexin-treated *Caulobacter crescentus* cell.

**Movie S2:** Tomogram of A22-treated *Caulobacter crescentus* cell.

**Movie S3:** Tomogram of A22-treated *Caulobacter crescentus* cell with envelope folding.

## METHODS

### SpyCatcher purification

His-tagged SpyCatcher conjugates were purified as previously described using nickel-affinity chromatography^31^. Plasmids pDEST14-SpyCatcher-sfGFP and pBAD-SpyCatcher-mRFP1 were transformed into chemical competent cells *E. coli* BL21 (DE3) and LMG194 cells respectively and grown on LB agar with 100 µg/mL Ampicillin (LB-Amp). A single colony of each strain was inoculated into 6 L of LB-Amp media and incubated at 37 °C with shaking until cells had reached mid-log growth phase. Cells were induced with 0.2% (w/v) arabinose (LMG194) or 0.4 mM Isopropyl β-D-1-thiogalactopyranoside (IPTG) (BL21) and incubated at 20 °C for 16 h. Induced cultures were harvested by centrifugation (15,000 relative centrifugal force (rcf), 30 mins, 4 °C), resuspended in lysis buffer (30 mM Tris/HCl pH 8.0, 500 mM NaCl, 1 mM MgCl_2_, 50 µg/mL DNase, 300 µg/mL lysozyme, and 1x cOmplete Protease Inhibitor), and lysed by five passes through the homogeniser at 15,000 psi (pounds per square inch) pressure. Cell debris were pelleted (50,000 (rcf), 45 mins, 4 °C), and the supernatant filtered using a 0.22 µm syringe filter. SpyCatcher proteins were then bound to a 5 mL HisTrap HP column (GE Healthcare) using an ÄKTA pure 25 M system (GE Healthcare) and eluted against the same buffer including 500 mM imidazole over 10 column volumes. Eluates were dialysed overnight with 1:100 (w/w) His_6_-TEV protease at 4 °C against 2 L of MilliQ H_2_O. The dialysates were further purified via size exclusion chromatography using a HiLoad Superdex S200 16/600 (prep grade) column; final proteins were eluted in HEPES buffer (25 mM HEPES/NaOH pH 7.5, 150 mM NaCl), and flash frozen in liquid nitrogen and stored at - 80 °C.

### SpyCatcher and HADA labelling of *C. crescentus*

*C. crescentus* expressing RsaA-467-SpyTag (CB15N ρ*sapA rsaA467:*SpyTag) cells were grown in PYE media (0.2% (w/v) Bacto Peptone, 0.1% (w/v) yeast extract, 0.5 mM CaCl_2_, 1 mM MgSO_4_) at 30 °C with aeration by shaking to mid-log growth phase. For SpyCatcher labelling, cells were resuspended to OD_600_ 0.1 in PYE, followed by pulse labelling with 10 µM SC-mRFP1 at 4 °C for 16 h, after which point cells were harvested by centrifugation (3 min, 8000 rcf) and washed three times with chilled PYE. For chase labelling, cells were resuspended in fresh PYE media and incubated at 30 °C for 1.5 h in the presence of 10 µM SC-sfGFP to stimulate growth. Samples treated with 50 µg/mL cephalexin or 3 µg/mL A22 were incubated with the SC-sfGFP chase for 3 h under the same conditions. After labelling, cells were harvested by centrifugation and washed as described above, followed by resuspension in a final volume of 50 µL PYE. For PG labelling, cells were supplemented with 500 µM of the fluorescent D-amino acid HADA^93^ (Cambridge Biosciences) for the last 10 minutes of the chase-labelling incubation. Cells were harvested and washed (as above). When required, cells were resuspended in chilled 4% formaldehyde (in PBS) for fixation. Samples were kept at 4°C for 20 minutes prior to washing (as above) and imaging. All incubation steps were carried out with the specimen protected from light-exposure. When necessary, cells were synchronised using density centrifugation method using colloidal silica^94^. Cells were grown and pulse labelled by incubation overnight with SC-mRFP1, as described. Cells were washed three times with PBS and resuspended in 750 µL ice-cold PBS. Samples were mixed 1:1 with syringe- filtered 33% chilled Percoll (Sigma-Aldrich), then centrifuged at 15,000 rcf in a tabletop centrifuge for 20 minutes at 4 °C, separating the cells into top (stalked cell) and bottom (swarmer cell) bands. The top band was carefully removed, and the bottom band collected, with a final volume 50-200 µL depending on the band size. Swarmer cell were pelleted and washed three times in ice-cold PBS media to remove excess Percoll. Cells were then resuspended in fresh PYE and chase labelled using SC-sfGFP as described above.

### Light microscopy

Two µL of labelled cell suspensions were spotted onto agarose pads (1% (w/v) in distilled water) enclosed by a 15 mm x 16 mm Gene Frame (ThermoFisher) on a glass slide and sealed with a glass coverslip. For cells labelled using only SpyCatcher conjugates, cells were imaged using an Olympus SoRa spinning disc confocal microscope, equipped with Olympus IX-83 inverted frame, Yokogawa SoRa super-resolution spinning disc module, and Prime BSI camera. Slides were kept at room temperature and imaged using the 60x (1.5 NA) lens, with excitation at 488 nm (sfGFP) and 561 nm (mRFP1) (solid state lasers), 200 ms exposure. Z-stacks were taken at 0.26 µm intervals and a super resolution filter applied to the entire stack using Olympus CellSens software, followed by deconvolution using a maximum likelihood algorithm (5 iterations). In general, Z-stacks were condensed using a maximum Z-projection of frames containing the cell of interest (± 1 frame on the upper and lower Z-axis). For HADA-labelled samples, cells were imaged using an Olympus Fluoview FV1200 equipped with equipped with GaAsP detectors. Images were acquired using a 100 x (1.4 NA) lens with excitation via solid state 405 nm (HADA) and 559 nm (mRFP1) and argon 488 nm (sfGFP) lasers, with scanning at 1024 x 1024 pixels. A Kalman filter (2 iterations) was applied for all image collections. Single slices through the middle of the cells were taken, without Z-stacks, to limit photobleaching of the sample. Images were background-subtracted and filtered using a 0.5-pixel Gaussian blur (ImageJ) unless stated otherwise.

### Light microscopy image analysis

Demographs and cell intensity profiles were generated using the MicrobeJ plugin for ImageJ^44^. Cell debris, overlapping cells, or cells on the edge of the micrographs were excluded from the analysis. The remaining cells were normalised for fluorescence intensity and plotted according to length from shortest to longest. Cell intensity profiles are representative of 100 dividing and non-dividing cells, assigned by the presence of invagination at the mid-cell, from the demograph. Normalised profile intensities from the sfGFP and mRFP1 channels, including standard deviation, were plotted relative to the cell length. For individual cell-profile analyses, a line was manually drawn along the indicated region of the cell through the cell body, straightened, and the pixel values extracted. Intensity values were normalised for each channel and plotted relative to the cell contour. Where given, cell profiles were binarized according to the presence of the strongest fluorescence intensity value. For colocalisation studies, masks were created for individual cells in ImageJ and colocalisation measured using Pearson’s Colocalisation Coefficient (PCC) in the Coloc2 plugin. A mask was created for individual cells and the colocalisation of the RFP1- and sfGFP-labelled regions was measured by PCC.

### Electron cryotomography (cryo-ET) Sample Preparation, Data Collection and Analysis

Cryo-ET grid preparation was performed as described previously^31, 64, 95^. Briefly, 2.5 µL of the relevant *C. crescentus* cell sample (OD_600_ 0.5-0.7 in PYE or M2G) mixed with 10 nm protein-A gold (CMC Utrecht) was applied to a freshly glow discharged Quantifoil R2/2 or R3.5/1Cu/Rh 200 mesh grid, adsorbed for 10 s, blotted for 2.5 s and plunge-frozen into liquid ethane in a Vitrobot Mark IV (ThermoFisher), while the blotting chamber was maintained at 100% humidity at 10 °C. For tomographic data collection, the SerialEM software^96^ was used as described previously^92^, using the Quantum energy filter (slit width 20 eV) and the K2 or K3 direct electron detector running in counting mode. Tilt series with a defocus range of -5 to -8 µm were collected between ± 65° in a bidirectional (only for Fig. 5A-C) or ±60° dose symmetric scheme with a 1° tilt increment. A total dose of 150 e^-^/Å^2^ (only Fig. 5A-C) or 73 e^-^/Å^2^ was applied over the entire series. Cryo-ET data analysis was performed in IMOD^97^ and tomographic reconstruction was carried out using the SIRT algorithm implemented within Tomo3D^98, 99^.

## ACKNOWLEDGEMENTS

M.H. was supported by funding from the Biotechnology and Biological Sciences Research Council (BBSRC, grant number BB/M011224/1). T.A.M.B. was a recipient of a Sir Henry Dale Fellowship, jointly funded by the Wellcome Trust and the Royal Society (202231/Z/16/Z). This work was supported by the Medical Research Council, as part of United Kingdom Research and Innovation (also known as UK Research and Innovation) [Programme MC_UP_1201/31]. For the purpose of open access, the MRC Laboratory of Molecular Biology has applied a CC BY public copyright licence to any Author Accepted Manuscript version arising. T.A.M.B. would like to thank the Human Frontier Science Program (Grant RGY0074/2021), the Vallee Research Foundation, the European Molecular Biology Organization, the Leverhulme Trust, and the Lister Institute for Preventative Medicine for support. We thank Jan Löwe, Buzz Baum and Buse Isbilir for critically reading this manuscript.

## MATERIALS AND CORRESPONDENCE

Requests should be addressed to Tanmay A. M. Bharat

